# An accelerometer-derived ballistocardiogram method for detecting heartrates in free-ranging marine mammals

**DOI:** 10.1101/2021.12.03.471142

**Authors:** Max F. Czapanskiy, Paul J. Ponganis, James A. Fahlbusch, T. L. Schmitt, Jeremy A. Goldbogen

## Abstract

Physio-logging methods, which use animal-borne devices to record physiological variables, are entering a new era driven by advances in sensor development. However, existing datasets collected with traditional bio-loggers, such as accelerometers, still contain untapped eco-physiological information. Here we present a computational method for extracting heartrate from high-resolution accelerometer data using a ballistocardiogram. We validated our method with simultaneous accelerometer-electrocardiogram tag deployments in a controlled setting on a killer whale (*Orcinus orca*) and demonstrate the method recovers previously observed cardiovascular patterns in a blue whale (*Balaenoptera musculus*), including the magnitude of apneic bradycardia and increase in heart rate prior to and during ascent. Our ballistocardiogram method may be applied to mine heart rates from previously collected accelerometery and expand our understanding of comparative cardiovascular physiology.

## Introduction

Recent advances in physio-logging (recording physiological variables using animal-borne devices) have largely been driven by new developments in sensor technology (Hawkes et al., 2021; Williams and Hindle, 2021). For example, new physio-logging tags can detect regional changes in blood flow by incorporating functional near-infrared spectroscopy sensors (McKnight et al., 2021). However, traditional inertial measurement unit (IMU) tags equipped with accelerometers and other inertial sensors can also measure important physiological and related variables. Through careful inspection and analysis of high-resolution acceleration, scientists have measured elevated respiration rates following record-breaking dives (Sato et al., 2011), near-continuous feeding by small cetaceans (Wisniewska et al., 2016), social interactions between large cetaceans (Goldbogen et al., 2014), and important biomechanical variables including movement speed (Cade et al., 2018). While physio-logging tags with cutting-edge biomedical technologies push the boundaries of physiological field research, simpler IMU tags have fewer logistical constraints and provide access to more species and larger sample sizes. This is particularly important for species that cannot be restrained or studied in managed care. For example, of the sixteen species of baleen whales (Mysticeti), heart rate has only been recorded with an electrocardiogram tag in the wild for one blue whale (*Balaenoptera musculus*) (Goldbogen et al., 2019; but see Ponganis and Kooyman, 1999). Conversely, IMU tags have been deployed on hundreds of individuals of nearly every species in the clade for the last twenty years (Nowacek et al., 2001). These existing datasets (and future IMU tag deployments) could hold additional valuable physiological information, awaiting proper computational methods for mining them.

The ballistocardiogram (BCG) has potential applications to using accelerometers as heartrate monitors in both the wild and in managed care. Ballistocardiography is a noninvasive method for measuring cardiac function based on the ballistic forces involved in the heart ejecting blood into the major vessels. The BCG originated as a clinical tool in the first half of the 20th century (Starr et al., 1939), but was largely superseded by electro- and echocardiography. However, potential novel applications like passive monitoring of heart function in at-risk populations (Giovangrandi et al., 2011) has led to a recent resurgence of ballistocardiography research, with advances in hardware (Andreozzi et al., 2021) and signal processing methodology (Sadek et al., 2019). While the BCG is a three-dimensional phenomenon, it is strongest in the cranio-caudal axis (Inan et al., 2015). Along this axis, the waveform is composed of multiple peaks and valleys; most prominent of these are the IJK complex, which progressively occurs during systole (Pinheiro et al., 2010). The BCG J wave is the most robust feature in the waveform and typically used for detecting heart beats.

Here we present a method for generating a BCG from bio-logger accelerometry. We validated our method with a simultaneously recorded electrocardiogram (ECG) on an adult killer whale in managed care (*Orcinus orca*) and applied it to detect heartrate in a blue whale. The relative orientation of the tag on the body is uncertain in cetacean bio-logging in the wild (Johnson and Tyack, 2003), so in addition to a one-dimensional BCG based solely on cranio-caudal acceleration, we also generated a three-dimensional BCG, which we expected would be more robust in a field setting. Specifically, we tested three hypotheses to validate our method. First, a one-dimensional BCG would, in a controlled setting, produce statistically equivalent instantaneous heartrates as an ECG. Second, a three-dimensional BCG would, in a field setting, produce a more robust signal than a one-dimensional BCG. Third, BCG-derived heartrates would increase during the latter phases of dives, consistent with the progressive increase in heartrate routinely observed prior to and during ascent (Goldbogen et al., 2019; McDonald and Ponganis, 2014).

## Materials and methods

### Animal tagging

#### Killer whale

A 3868 kg adult female killer whale in managed care at SeaWorld of California, San Diego, CA was double-tagged with Customized Animal Tracking Solutions IMU (CATS, www.cats.is) and electrocardiogram (ECG) tags on August 16, 2021 as part of clinical animal cardiac evaluations under the SeaWorld USDA APHIS display permit. We attached the CATS tag on the mid-lateral left chest posterior to the pectoral fin (Movie S1). The CATS tag recorded acceleration at 400 Hz, magnetometer and gyroscope at 50 Hz, pressure at 10 Hz, and video at 30 fps. All sensors were rotated from the tag’s frame of reference to that of the whale using MATLAB (MathWorks, Inc., v2020b) tools for processing CATS data (Cade et al., 2021). This rotation aligned the tag’s x-, y-, and z-axes with the cranio-caudal, lateral, and dorso-ventral axes of the whale, respectively. The ECG tag hardware and data processing followed the methods in (Bickett et al., 2019). Briefly, the tag was attached approximately midline on the ventral chest just caudal (posterior) to the axilla and we recorded the ECG at 100 Hz. Individual heart beats were identified from visually verified R-waves using a customized peak detection program (K. Ponganis; Origin 2017, OriginLab Co., Northampton, MA). ECG and IMU were recorded during a spontaneous breath hold while the whale rested at the surface.

#### Blue whale

A 24.5 m blue whale was tagged with a CATS IMU tag on September 5, 2018 in Monterey Bay, CA under permits MBNMS-MULTI-2017-007, NMFS 21678, and Stanford University IACUC 30123 (previously published by Gough et al., 2019). The tag slid behind the left pectoral flipper, similar to the placement of the CATS tag on the killer whale. Tag configuration and data processing followed the same procedure as the killer whale. The 400 Hz acceleration data was used for ballistocardiography (see section **Signal processing**). We downsampled the multi-sensor data to 10 Hz for movement analysis using the MATLAB CATS tools.

### Signal processing

The BCG waveform is three dimensional, but strongest in the cranio-caudal axis (Inan et al., 2015). We tested both 1-dimensional (cranio-caudal only) and 3-dimensional metrics for identifying heartbeats in acceleration data based on the methods of (Lee et al., 2016). For windowed operations, we used 0.5 s for killer whale data and 2.0 s for blue whale data.

#### Procedure

1. Remove noise and de-trend the acceleration signal with a 5th order Butterworth band-pass filter (killer whale: [1-25Hz], blue whale: [1-10Hz]) (R package signal) (Ligges et al., 2021).
2. Enhance the IJK complex by differentiating acceleration using a 4th order Savitzky-Golay filter (R package signal). Differentiation exaggerates impulses like the J wave.
3. Further enhance the peaks by calculating the Shannon entropy (*H*_*i*_ = − ∑ *k* | *a*_*i*_*k* | × *In*(|*a*_*i*_*k* |), where *k* is the acceleration axis). Additionally, the Shannon entropy is strictly positive, which facilitates peak detection. In the 1-dimensional case, *k* is surge (cranio-caudal acceleration) only.
4. Remove noise by applying a triangular moving average smoother.
5. Extract peaks and heuristically remove noisy peaks (Fig. S1).

This procedure may be applied to either 1-dimensional (i.e., cranio-caudal only) or 3-dimensional acceleration. In the case of 3-dimensional acceleration, the band-pass and Savitzky-Golay filters were applied to each axis independently.

### BCG validation with killer whale ECG

We fit ordinary least squares regression to BCG-derived instantaneous heart rates with respect to ECG-derived and tested 1) if the intercept was significantly different than 0 and 2) if the slope was significantly different than 1. We calculated the mean and standard deviation of absolute error as an equivalence measure (1-dimensional BCG only).

### BCG application to blue whale

Dynamic body movements produce an acceleration signal that masks the ballistocardiogram, so we limited our analyses to motionless periods (Fig. S2). These periods occured during or near the bottom phase of dives between fluke strokes. Strokes were detected from visual examination of the rotational velocity around the lateral axis recorded by gyroscope (*sensu* Gough et al., 2019). We tested whether the 3-dimensional BCG was more robust than 1-dimensional BCG in field data by comparing the signal-to-noise ratios. For both BCGs, we calculated the power spectral density (R package psd) (Barbour and Parker, 2014). Previously recorded blue whale apneic heart rate was 4-8 beats per minute (bpm) (Goldbogen et al., 2019), so we quantified *signal* as the integration of the power spectral density curve from 4-8 bpm and *noise* as the integrated remainder, up to 60 bpm.

We also tested whether BCG-derived instantaneous heart rates were consistent with the range and pattern of heart rates previously observed in the blue whale and other marine mammals; namely a gradual increase in heart rate later in the dive, especially during the final ascent (Goldbogen et al., 2019; McDonald and Ponganis, 2014). We assigned dive start and end times when the whale swam deeper than 2 m, retaining dives that exceeded 10 m depth and 5 minutes duration. Dive times were normalized from 0 (start of dive) to 1 (end of dive). We regressed instantaneous heart rate against normalized dive time using robust Theil-Sen regression (to account for heteroscedascity) (R package RobustLinearReg) (Hurtado, 2020; Sen, 1968; Theil, 1992) and tested whether the slope was greater than 0.

### Reproducibility

The data and code used in this analysis were packaged as a research compendium (R package rrtools) (Marwick, 2019; Marwick et al., 2018). The research compendium was written as an R package so other researchers can read, run, and modify the methods described here.

## Results and discussion

### BCG validation with killer whale ECG

The ECG and BCG yielded nearly identical heart rate estimations (Fig. 1). We collected 14 s of simultaneous ECG and BCG data during a motionless breath hold at the surface. BCG-derived instantaneous heart rates were within 0.8% ± 0.5% of the ECG-derived rates (mean ± standard deviation). Ordinary least squares regression of BCG heartrates on ECG heartrates yielded a slope of 1.02 ± 0.04 and intercept of −1.62 ± 2.71 (mean ± standard error), which were not significantly different from the hypothesized 1 and 0, respectively.

**Figure 1:**
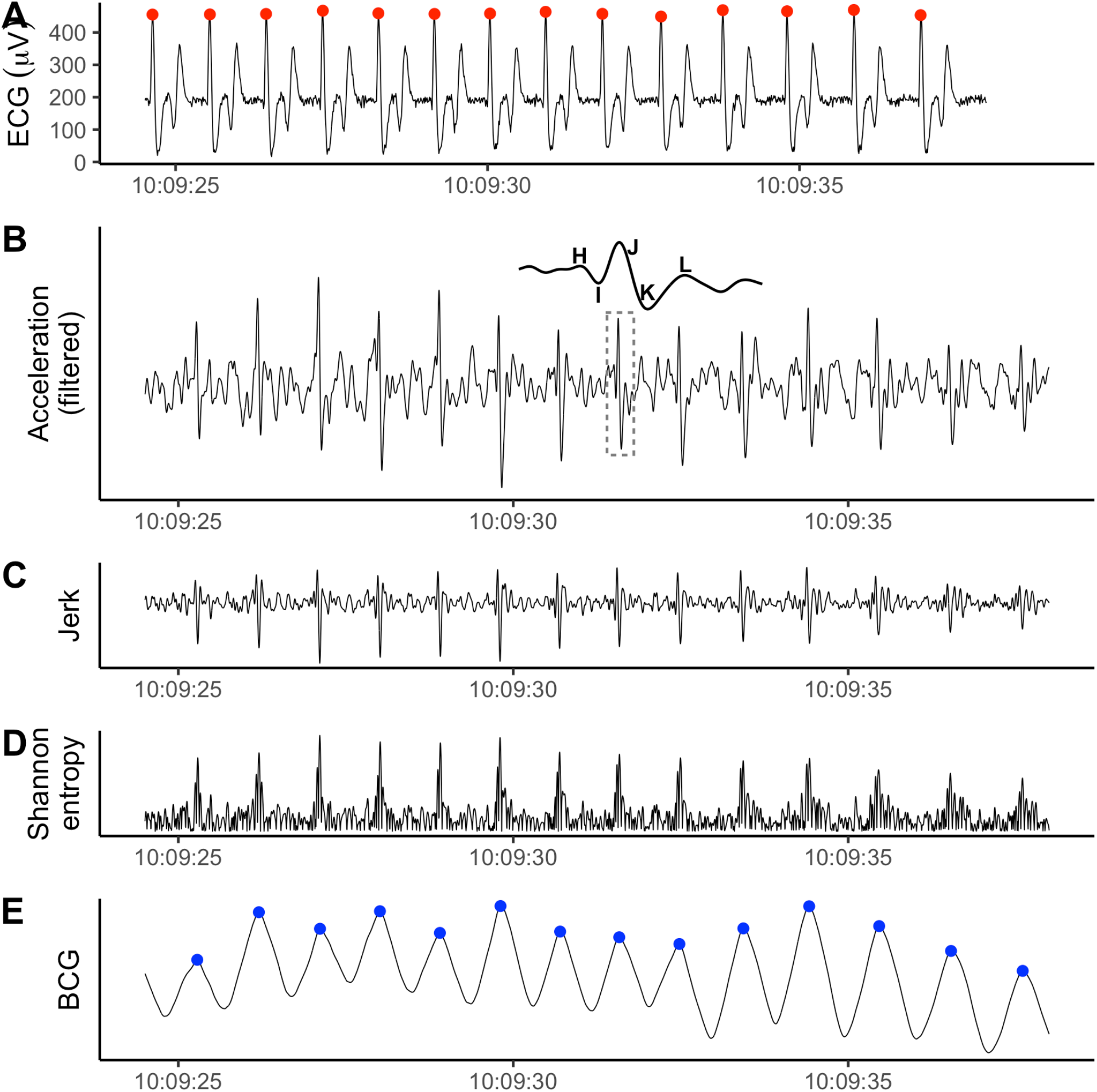
The ECG (**A**, recorded by ECG tag) and 1-dimensional BCG (**E**, processed from the cranio-caudal acceleration recorded by the IMU tag) produced nearly identical heart beat predictions for the killer whale. **B-D** display the intermediate steps in the BCG signal processing procedure. **B**: Cranio-caudal axis acceleration after band-pass filtering. Inset shows the IJK complex with surrounding H and L waves for the region bounded by the dashed box. **C**: Peaks enhanced after forward differencing acceleration (i.e., jerk). **D**: A strictly positive signal after calculating Shannon entropy. Y-axis values excluded because filtering introduces magnitude distortion and only the relative shape of the signal is relevant to the analysis.

### BCG application to blue whale

We generated 1-dimensional and 3-dimensional BCGs for 2 hours of data, including 10 rest dives and 51 motionless periods totaling 76.9 minutes (Fig. S3).

The 3-dimensional BCG (Fig. 2) produced a more robust signal than the 1-dimensional BCG, which used only cranio-caudal acceleration. The signal-to-noise ratio was 2.00 for the 3-dimensional BCG, compared to 0.17 for the 1-dimensional BCG (Fig. 3**A**).

**Figure 2:**
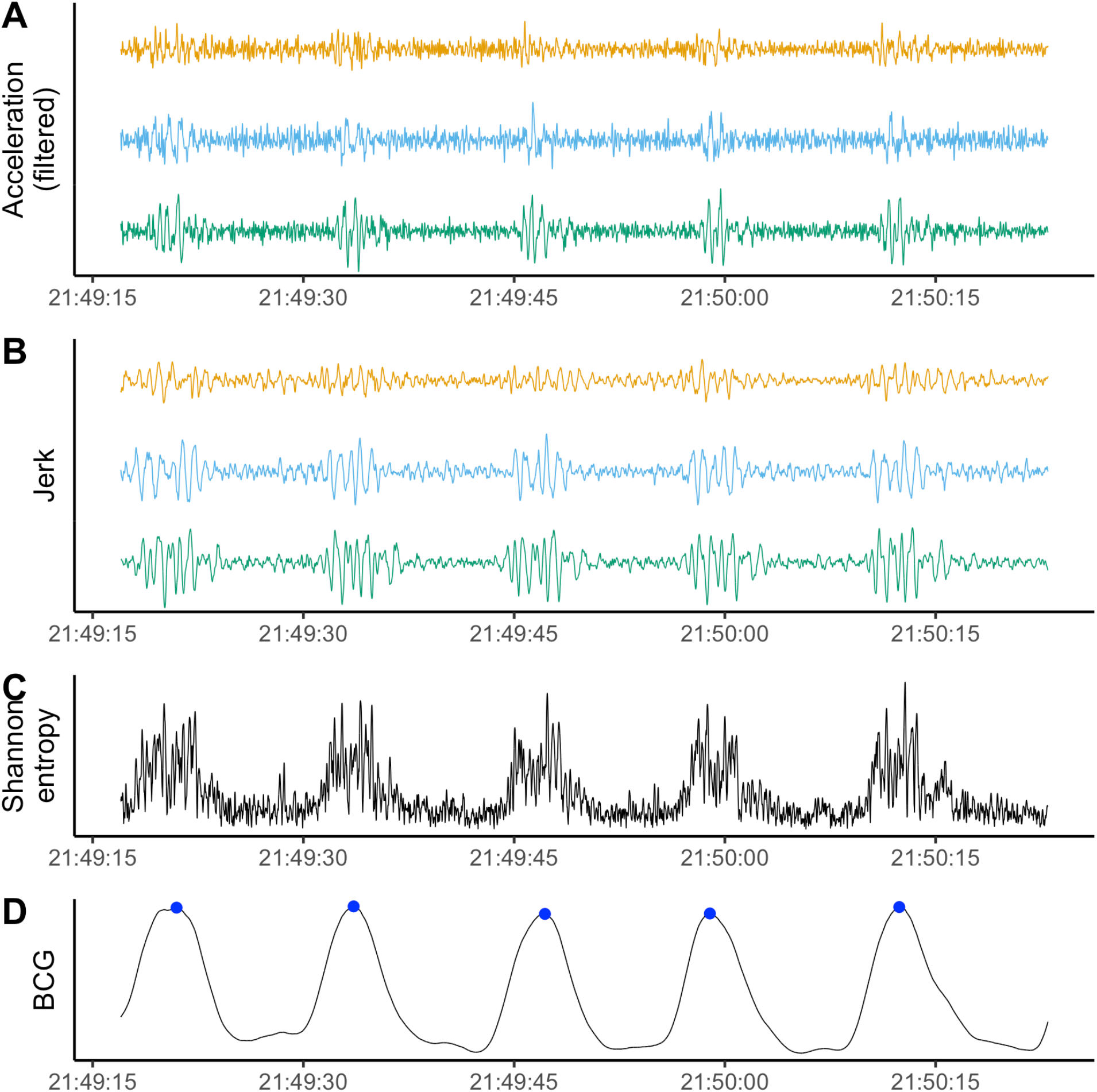
Example of signal processing for 3-dimensional BCG during a motionless period in a blue whale dive. **A**: Band-pass filtered triaxial acceleration, with cranio-caudal in orange, lateral in blue, and dorso-ventral in green. **B**: Peaks enhanced after forward differencing acceleration (i.e., jerk). **C**: The Shannon entropy combines information from all three axes and makes the signal strictly positive. **D**: Smoothing the Shannon entropy facilitates robust peak detection. Detected heart beats in blue. Y-axis values excluded because the filtering process introduces magnitude distortion and only the relative shape of the signal is relevant to the analysis.

**Figure 3:**
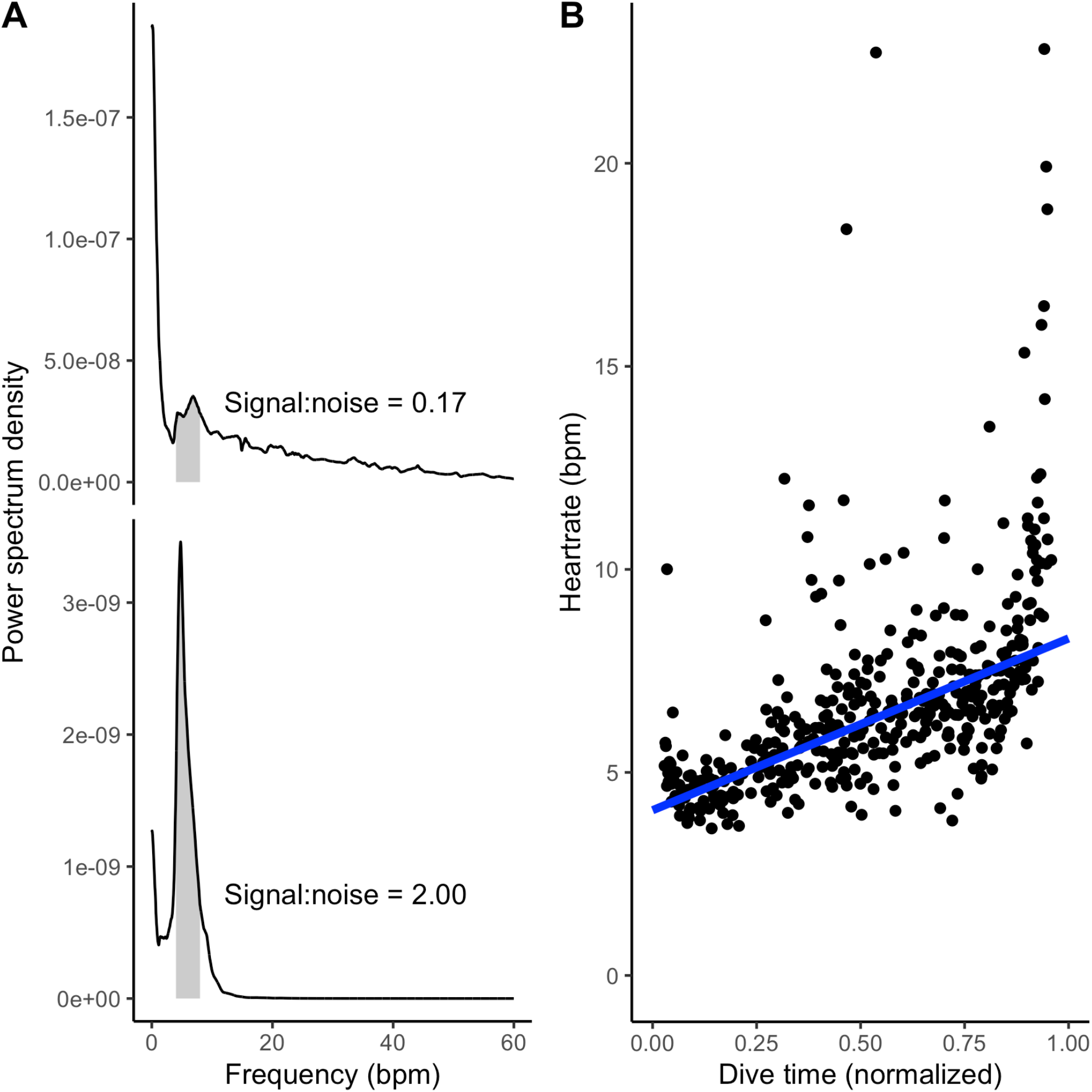
**A** Signal-to-noise ratio was higher for the 3-dimensional BCG (lower panel) than the 1-dimensional BCG (cranio-caudal acceleration only; upper panel). Each panel shows the power spectral density for the BCG. Based on previously observed blue whale heart rates, 4-8 bpm was considered signal (gray shading). The signal-to-noise ratio was calculated as the ratio of the area under the curve in the signal band to the area under the rest of the curve, up to 60 bpm. **B** Heart rates observed in the 3-dimensional BCG followed characteristic diving physiology patterns. Heart rate is lowest at the start of the dive (∼4-5 bpm), increasing towards ascent (∼8-9 bpm). Points indicate instantaneous heart rates and the line is a Theil-Sen regression. Outliers likely represent premature beats which are common in heart rate profiles during dives of cetaceans, pinnipeds, and penguins (Andrews et al., 1997; Goldbogen et al., 2019; McDonald and Ponganis, 2014; Wright et al., 2014).

3-dimensional BCG-derived heart rates exhibited a relaxation of bradycardia over the course of dives. Average heart rate increased from 4.1 bpm at the start of dives to 8.3 bpm at the end of dives (Theil-Sen regression, *p*< 10 ^-10^(Fig. 3**B**).

### Reproducibility

The research compendium containing data, code, and an executable version of this manuscript was archived on Zenodo (Czapanskiy, 2021). We developed the research compendium as an R package to facilitate investigation and adoption by other researchers. Publishing data and code in standardized formats (such as an R package) is a critical step towards transparency and computational reproducibility (Alston and Rick, 2021; Powers and Hampton, 2019; Stodden et al., 2018).

## Conclusions

Here we presented a ballistocardiogram method for detecting resting apneic heartrate in cetaceans using accelerometers. We validated the method in a controlled setting with simultaneous ECG and in a field setting by confirming expected physiological patterns. As accelerometer tags have been deployed on many cetacean species for multiple decades, this method may be applied to mine existing datasets and better understand how heartrate scales with body size and other biological factors. Current IMU tag designs limit BCG analysis to motionless periods, but future dimple-or limpet-style tags could reduce acceleration noise, boost the signal-to-noise ratio, and make the method more widely applicable. Even as the field of physio-logging progresses with new hardware innovations, this method demonstrates that computational advances can still derive new insights from traditional sensors.

## Acknowledgements

The authors are grateful to the SeaWorld of California Killer Whale training staff for their efforts and support. We also thank Anna Krystalli, Ben Marwick, Karthik Ram, Nicholas Tierney, and other members of the R community for developing tools and educational resources to facilitate open science practices. This is a SeaWorld Parks and Entertainment Technical Contribution number 2021-12.

## Footnotes

## Author contributions

Conceptualization: M.F.C.,J.A.F.,P.J.P.,J.A.G.; Methodology: M.F.C.,J.A.F.,P.J.P.; Software:

M.F.C.; Formal analysis: M.F.C.,P.J.P.; Investigation: M.F.C.,J.A.F.,P.J.P.; Resources: P.J.P.,J.A.G.; Writing - original draft: M.F.C; Writing - review & editing: M.F.C.,J.A.F.,P.J.P.,J.A.G.; Supervision: P.J.P.,J.A.G.; Project administration: P.J.P.,J.A.G.; Funding acquisition: J.A.G.

## Data availability

All data and code used in this analysis are available on Zenodo (DOI needed).

## Colophon

This report was generated on 2021-12-03 11:06:51 using the following computational environment and dependencies:

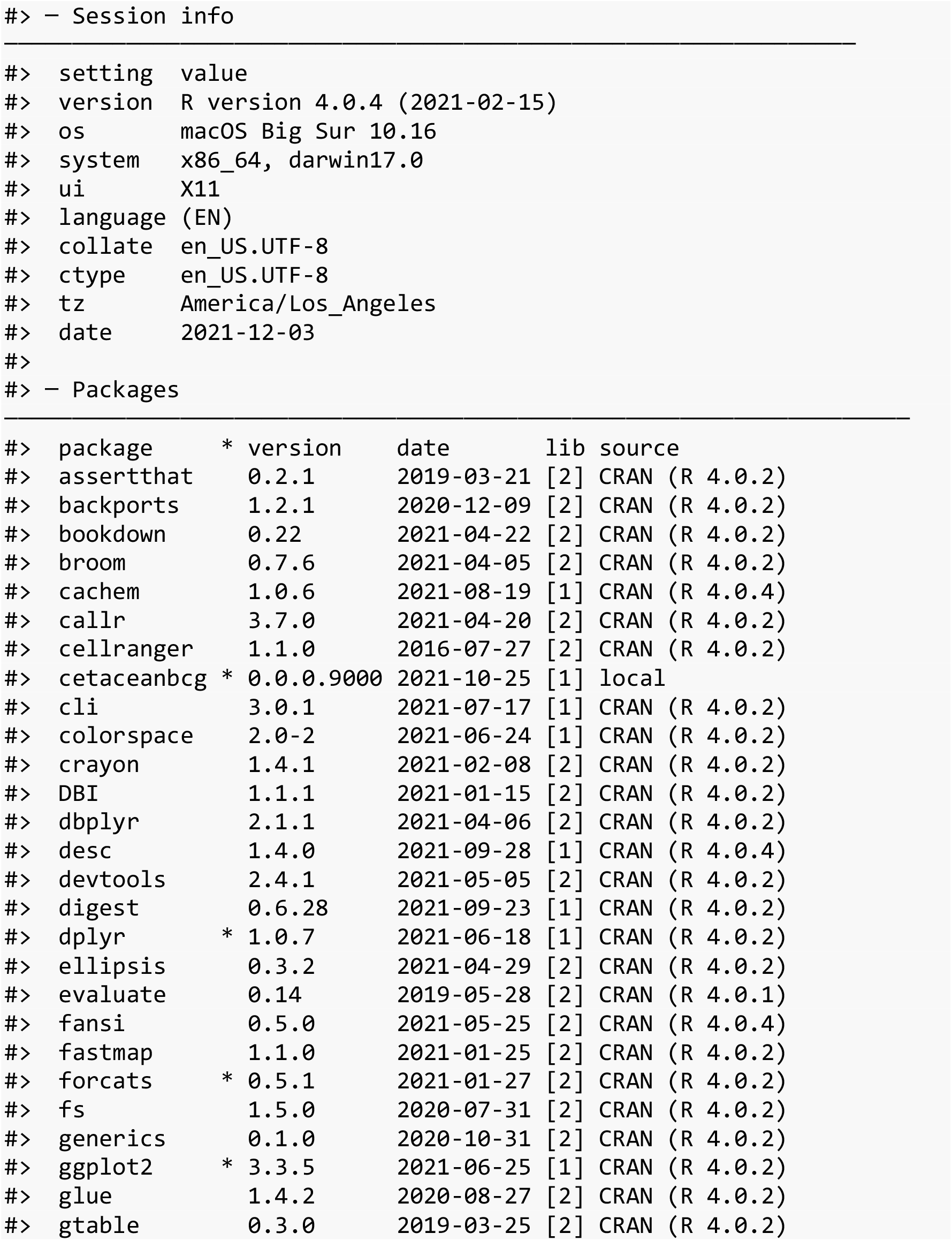
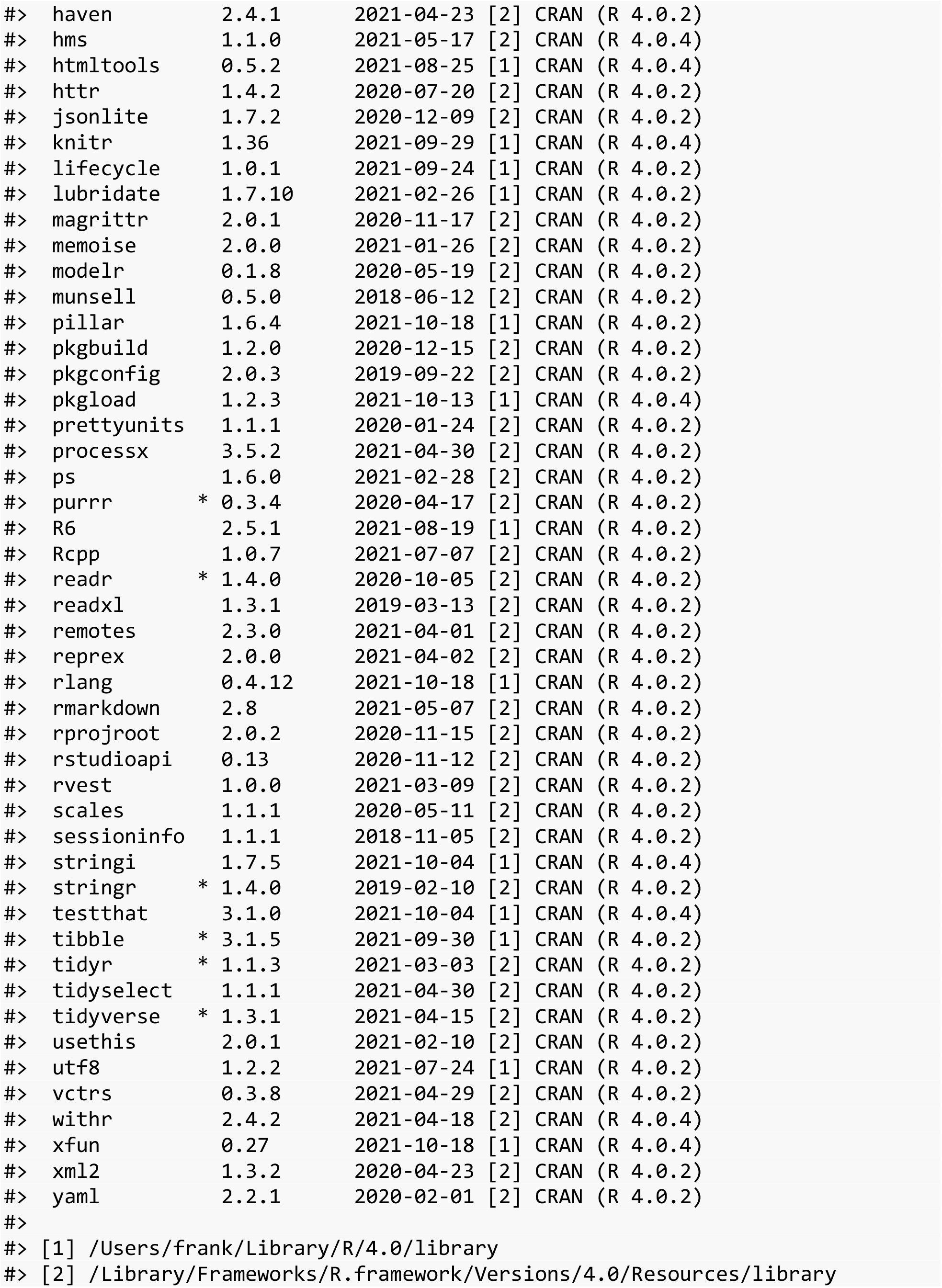

The current Git commit details are:

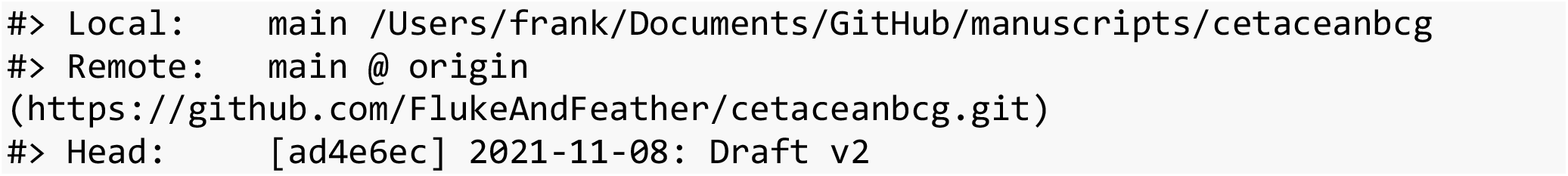

