## Supplemental figures for "An accelerometer-derived ballistocardiogram method for detecting heartrates in free-ranging marine mammals"

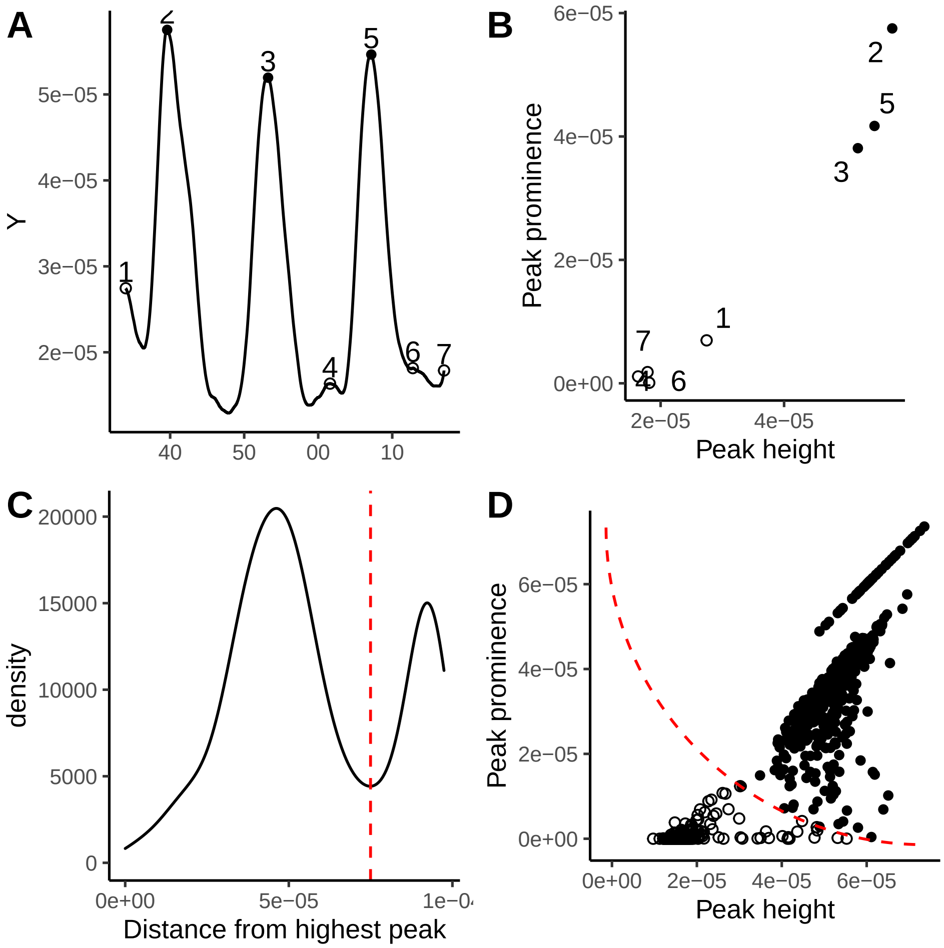


Figure S1: **A:** Minor peaks (hollow points) in the ballistocardiogram (BCG, line) were not considered heart beats. Only major peaks (solid points) were retained for analysis. The BCG for one motionless period shown here. **B:** We used peak height and prominence (i.e. height relative to the contour surrounding a higher peak) to heuristically differentiate major and minor peaks. For each peak, we calculated the Euclidean distance (in height-prominence space) to the highest peak overall. The peaks in **A** shown here in height-prominence space. **C:** The distance to the highest peak exhibited a bimodal distribution. We chose a distance threshold (dashed red line) corresponding to the valley in the density curve. **D:** All peaks found in the BCG across all motionless periods in height-prominence space. Solid and hollow points as in **A**. The dashed red curve corresponds to the distance threshold in **C**.

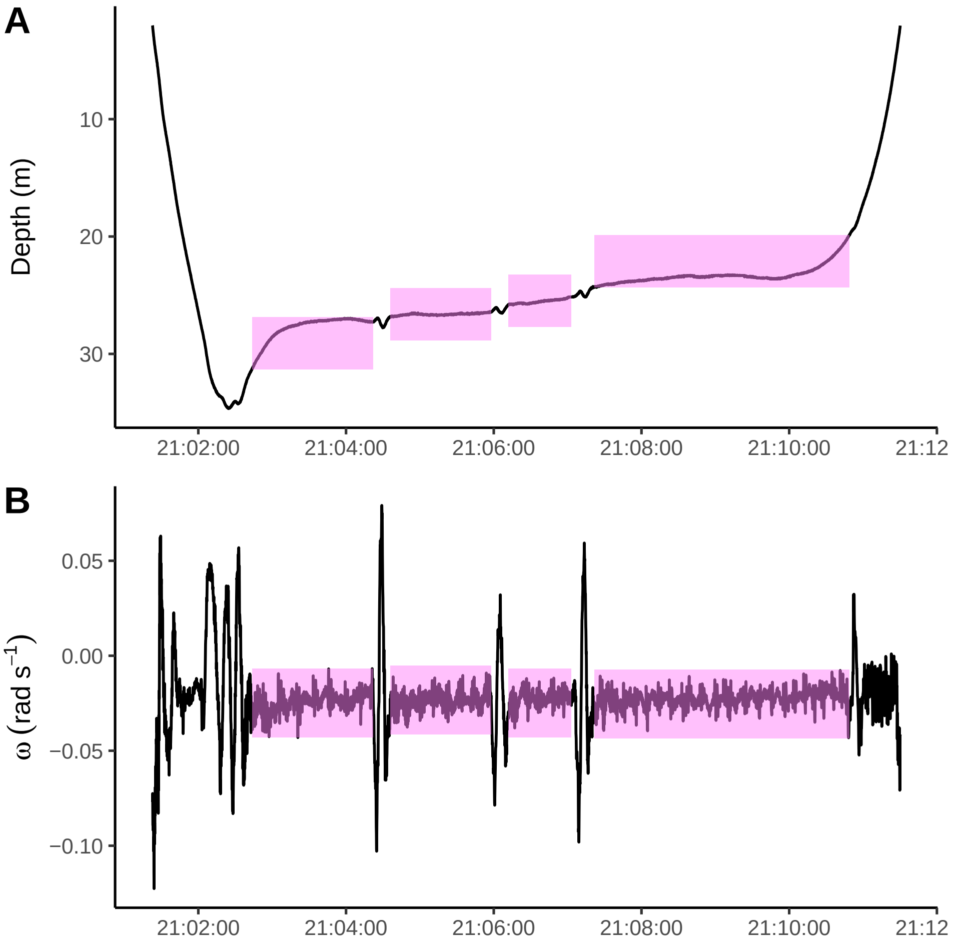


Figure S2: Motionless periods were manually identified by looking for periods of low amplitude rotational veloctiy ($\omega$) around the lateral axis, indicative of cetaceans' dorso-ventral fluking motion. The depth and $\omega$ profiles for the first dive shown here, with motionless periods in pink.

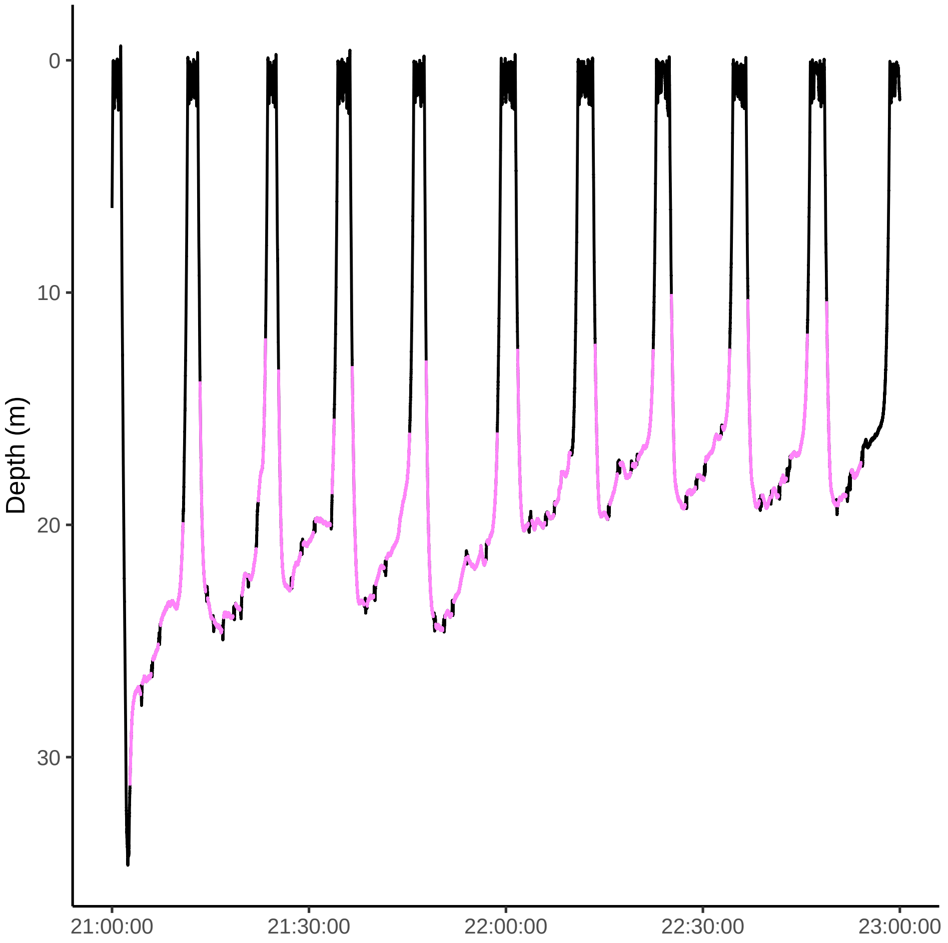


Figure S3: Two hours of blue whale tag data were used for ballistocardiographic analysis. During these hours, the animal made repeated, shallow dives. Because the BCG can be obscured by dynamic body movements, only relatively motionless periods were retained for analysis (pink). These periods were identified by visual examination of the rotational velocity around the lateral axis.
